# A screen for Twist-interacting proteins identifies Twinstar as a regulator of muscle development during embryogenesis

**DOI:** 10.1101/2021.02.10.430492

**Authors:** Mridula Balakrishnan, Austin Howard, Shannon F. Yu, Katie Sommer, Scott J. Nowak, Mary K. Baylies

**Affiliations:** Biochemistry & Structural Biology, Cell & Developmental Biology, and Molecular Biology (BCMB) Program, Weill Cornell Graduate School of Medical Sciences, New York, NY 10065; Department of Molecular and Cellular Biology, Kennesaw State University, Kennesaw, GA 30144; Master of Science in Integrative Biology Program, Kennesaw State University, Kennesaw, GA 30144; Developmental Biology Program, Sloan Kettering Institute, Memorial Sloan Kettering Cancer Center, 1275 York Avenue, New York, New York 10021

**Keywords:** Twist, bHLH, transcription, mesoderm, muscle

## Abstract

Myogenesis in *Drosophila* relies on the activity of the transcription factor Twist during several key events of mesoderm differentiation. To identify the mechanism(s) by which Twist establishes a unique gene expression profile in specific spatial and temporal locales, we employed a yeast-based double interaction screen to discover new Twist-interacting proteins (TIPs) at the *myocyte enhancer factor 2 (mef2)* and *tinman (tinB)* myogenic enhancers. We identified a number of proteins that interacted with Twist at one or both enhancers, and whose interactions with Twist and roles in muscle development were previously unknown. Through genetic interaction studies, we find that Twinstar (Tsr), and its regulators are required for muscle formation. Loss of function and null mutations in *tsr* and its regulators result in missing and/or misattached muscles. Our data suggest that the yeast double interaction screen is a worthy approach to investigate spatial-temporal mechanisms of transcriptional regulation in muscle and in other tissues.

## INTRODUCTION

Spatiotemporal regulation of gene expression is crucial to direct the myriad cellular processes that work in concert to pattern a developing organism. This is often accomplished via the regulation of transcription factor (TF) activity during development. One example of this phenomenon is the bHLH transcription factor, Twist, which can orchestrate unique transcription profiles during mesoderm specification, patterning, and differentiation throughout *Drosophila* embryogenesis (Baylies *et al*. 1998; Castanon and Baylies 2002). However, how Twist mediates these diverse transcriptional programs in a spatiotemporal manner remain unclear.

Twist is dynamically expressed in the mesodermal cell lineage during *Drosophila* development. Prior to gastrulation, Twist is expressed in mesoderm precursors, and its expression is maintained as the cells invaginate, proliferate, and migrate dorsally during germ band extension. Once gastrulation is complete, Twist expression is confined to a segmentally repeated pattern of high and low protein levels. These Twist protein levels are critical for the specification and differentiation of mesodermal cells into different tissue types: high Twist-expressing cells form the somatic or body wall muscles and heart, while low Twist levels permit cells to form other mesodermal tissues, including the fat body and visceral musculature (Baylies and Bate 1996; Wong *et al*. 2008; 2014). As somatic myogenesis proceeds, Twist is expressed transiently in subsets of muscle founder cells, governing their identity (Wong *et al*. 2008; 2014). At the end of embryogenesis and through the larval stages, Twist is maintained in a specialized set of founders, the adult muscle progenitors (AMPs). Here, Twist prevents the AMPs from prematurely differentiating into the adult musculature (Cripps *et al*. 1998). All these distinct cellular events are governed by specific and discrete levels of Twist activity, which correspond to Twist expression levels (Wong *et al*. 2014). These Twist-regulated processes are highly sensitive to slight perturbations in Twist activity levels (Wong *et al*. 2014). How Twist’s activity, levels, and its mechanism of regulating varied transcriptional programs remain unclear.

A well-known mechanism through which Twist activity is modulated during both early and late mesodermal patterning is through its interactions with dimerization partners, including itself or Daughterless (Castanon *et al*. 2001), (Wong *et al*. 2008), as well as secondary gene-binding proteins such as Dorsal (González-Crespo and Levine 1993; Shirokawa and Courey 1997; Pham *et al*. 1999). While the importance of interactions with these partners for modulating Twist outputs has been demonstrated for myogenesis, they fail to account for the numerous developmental contexts that rely upon Twist during mesodermal specification and differentiation. This argues for the existence of other Twist-interacting proteins (TIPs), that function along with Twist throughout myogenesis or at discrete steps during mesoderm development. Therefore, identification of these TIPs is critical to understand the morphogenetic events that are required to form a functional muscle fiber.

Here we report the identification of 37 novel mesodermally-expressed Twist-interacting proteins (TIPs) through a yeast-based double interaction screen. This screen was designed to recover proteins that interact with Twist on one or both of two Twist-regulated myogenic enhancers. The first enhancer, the Twist-dependent *Dmef2* enhancer is critical for *Dmef2* expression, which governs the formation of the body wall musculature (Cripps *et al*. 1998; Nguyen and Xu 1998; Cripps *et al*. 1999). The second enhancer, the Twist-dependent *tinman* enhancer (*tin*B), is essential for establishing the cardiac myogenic program during embryogenesis (Yin *et al*. 1997; Yin and Frasch 1998). Upon analyzing the TIPs identified through our screen, we discover a new role for Twinstar (Tsr), an actin depolymerizing factor, in embryonic muscle development and patterning. Our data suggest that our screen is a powerful technique that can be used to identify new co-regulators of TFs during development.

## MATERIALS AND METHODS

### Double Interaction Screen in *S. cerevisiae*

#### enhancer-reporter yeast constructs

The plasmid 211-HIS was constructed by cloning a *XbaI/PstI* fragment of HIS3 from pUC19 HIS(D/X) (Yu *et al*. 1999) into YIplac211(Gietz and Akio 1988). The Twist-regulated *Dmef2* enhancer (Cripps *et al*. 1999) was cloned upstream of HIS in the 211-HIS plasmid using the 5’ primer CGCGAATTCTGGAGATGAGTTTCACGTGG (*Eco*RI end) and the 3’ primer CGCTCTAGATGTGCGCCGTACGGTTG (*Xba*I end) to PCR *Drosophila melanogaster* genomic DNA. The *tinman* enhancer (Yin *et al*. 1997) was cloned upstream of HIS in the 211-HIS plasmid using the 5’ primer CGCGAATTCCTCGAGGCTTTGACAAATCATC (EcoRI end) and the 3’ primer CGCTCTAGAGCGGGAAATGGAAAAGCG (XbaI end) to PCR *D. melanogaster* genomic DNA. These constructs were confirmed by sequencing. The *Dmef2* and *tinman* constructs were integrated into the *URA* locus of the yeast strain *NLY2* (*MATα gal4Δ gal80Δ his3 lys2 ura3-52 leu2 trp1*), creating two new strains *Dmef2/HIS* and *tinman/HIS* respectively.

#### twist constructs

A low copy vector was designed to express Twist in the following manner: first, the *Hind*III site of YCplac22, described in (Gietz and Akio 1988) was eliminated through digestion, a Klenow reaction and ligation (YCplac22H3KO). The ADH promoter and terminator were taken from the pADNS vector by digestion with BamHI and cloned into YCplac22H3KO, creating the vector 22-ADH. Twist was cloned into the HindIII and NotI sites of the multiple cloning region in the ADH region, creating 22-Twist. 22-Twist was transformed into the *Dmef2/HIS* and *tinman/HIS* strains. Prior to performing the screen, the optimal level of 3-AT concentration was determined by transforming this strain with the empty vector, pACT2 and plating concentrations of 3-AT in increments of 2.5mM from 0 to 15mM on plates lacking histidine.

#### Screen

A library scale transformation was performed on each of *Dmef2-HIS + 22-Twist* and *tinman-HIS + 22-Twist* strains using a 0-6 hr *Drosophila* embryonic library (a gift of L. Pick) according to the manufacturer’s instructions (Clontech PT3024-1). Transformations were plated on 150mm plates containing 12.5mM 3-AT. A control containing the empty vector (pACT2) was compared to the experimental condition. Lysates from positive colonies were transformed directly into *E. coli*. To determine plasmid dependence, plasmids isolated from *E. coli*were transformed back into the yeast parental strain from which the positive clone had been isolated. Only transformants that retested positive were kept. Positive cDNAs were sequenced and BLAST searches performed against the fly genome to determine cDNA identity. Obvious false positives, such as ribosomal and mitochondrial proteins, were eliminated at this stage (Serebriiskii *et al*. 2000).

### Secondary screening in Yeast

#### Twist dependence

Twist dependence was determined by transforming each of the cDNAs into their respective parental strains lacking Twist (*Dmef2/HIS* and *tinman/HIS*). Plasmids that failed induce growth, due to their inability to activate histidine synthesis, would, in this screen, be dependent on Twist to activate transcription, while those that were still able to activate HIS3 would do so in a Twist-independent manner.

#### Enhancer specificity

Positive cDNAs recovered in one enhancer screen were also tested on the other enhancer to determine whether the isolated cDNA was enhancer specific.

#### Mapping of Twist activity domains

Twist deletion constructs were generated via PCR using the following primer pairs. Twist (full length construct) Forward: 5’ CATG<u>CCATGG</u>AAATGATGAGCGCTCGCT-3′ and Reverse: 5′-G<u>GAATTC</u>CCTGATCCGCCGCTATG-3′. 143-490 (Δ AD2 domain) Forward: 5′-CATG<u>CCATGG</u>CATCCTCTTGGAACGAGCACGGCA-3′ and Reverse: 5′-G<u>GAATTC</u>CCTGATCCGCCGCTATG-3′. 1-331 (Δ bHLH and ΔWR motifs) Forward: 5′-CATG<u>CCATGG</u>AAATGATGAGCGCTCGCT-3′ and Reverse: 5′-G<u>GAATTC</u>CGTCCA GCAAACTGCCGGCACT-3′. 1-468 (ΔWR motif) Forward: 5′-CATG<u>CCATGG</u>AAATG ATGAGCGCTCGCT-3′ and Reverse: 5′-G<u>GAATTC</u>GGGCGGGATAATGGGTGCT-3′. 1-420 (Δ C-terminus and ΔWR motif) Forward: 5′-CATG<u>CCATGG</u>AAATGATGAGCG CTCGCT-3′ and Reverse: 5′-G<u>GAATTC</u>CCTGATCCGCCGCTATG-3′. Underlined sequences correspond to either *BamH*I or *N*coI sequences added to facilitate cloning of PCR product. Constructs were subsequently cloned into the *Hind*III and *Not*I sites of the 22-ADH vector (Mumberg *et al*. 1995). The 1-420 construct was created through digestion of wildtype Twist cDNA with *Xba*I, which results in the 3’ deletion of Twist at amino acid 420. Deletion of Twist activation domain 1 (ΔAD1 construct), consisting of amino acids 1-141 fused in-frame to amino acids 330-490 was originally constructed by digesting Twist with *Bam*HI, releasing a fragment containing the amino acids 142-329. Ligating the remaining fragments resulted in an in-frame deletion of these amino acids.

All constructs tested, as well as wildtype Twist were cloned into the *HindIII* and *NotI* sites of the ADH region of 22-ADH. Constructs were sequenced to ensure accuracy.

### *Drosophila* stocks

Unless otherwise noted, *Drosophila melanogaster* stocks were obtained from the Bloomington, Szeged, and/or Exelixis stock centers. *sina^2^* flies were a gift from R. Carthew. Second chromosome mutants were crossed to *Kr^If^CTG (CyO,P[GAL4-twi.G]2.2,P[UAS-2xEGFP]AH2.2)* flies and third chromosome mutants to *D*gl^3^/TTG (TM3,P[GAL4-twi.G]2.3, P[UAS-2xEGFP]AH2.3,Sb^1^Ser^1^)* flies in a balancer exchange to create new stocks carrying a fluorescent balancer. The *twist* null mutant stock *twi^1^/CyO, P[ry^+t7.2^=en1]wg^en11^* was used for examining double heterozygous flies. *UAS-2XeGFP* (Halfon *et al*. 2000), *UAS-moesin::GFP* and *UAS-moesin::mCherry* (gifts of J. Zallen), *apME-NLS::dsRed* (Richardson *et al*. 2007) *twist-Gal4* (Baylies *et al*. 1995), and *Dmef2-Gal4* (a gift from A. Michelson) were also used. *OreR* and *yw* were used as wild-type strains. Stocks used were *tsr^N96A^* (BDSC #9108), *tsr^1^* (BDSC #9107), *flr^1^* (BDSC #1132), *flr*^3^ (BDSC #2371), *ssh^1-63^* (BDSC #9110), *ssh^1-11^* (BDSC #9111), *LIMK1^EY08757^* (BDSC #17491), *cdi^07013^* (BDSC #11711), and *Mhc-Gal4* (BDSC #67044). The following gene traps were used: *tsr::GFP (ZCL2393)* (DGRC #110875) and *flr::GFP* (CA07499) (BDSC #50824).

### Immunohistochemistry

Embryos were collected at 25°C (unless indicated otherwise) on either apple or grape juice agar plates and were fixed in 4% paraformaldehyde/heptane. Embryos were mounted in Prolong Gold (Molecular Probes) or Vectashield (Vector Labs). Antibodies were used at the indicated final dilutions. Antibodies used were rat anti-Tropomyosin (1:1000, Abcam), mouse anti-Myosin heavy chain (1:500, gift from S. Abmayr), chicken anti-β-Galactosidase (1:1000, Abcam), rabbit anti-dsRed (1:400, Clontech), and mouse anti-GFP (1:200; PA; Clontech). Alexa Fluor 488-, Alexa Fluor 555-, and Alexa Fluor 647-conjugated secondary antibodies were used at 1:400 (Invitrogen).

For imaging of protein traps, embryos were collected and dechorionated as above and mounted in Halocarbon 700 Oil (Halocarbon Products) on a glass slide and covered with a coverslip. Slides were kept overnight and imaged the following day. Fluorescent confocal images were acquired on either a Zeiss LSM 700 confocal microscope equipped with a Plan-Apochromat 20x/0.8 WD=0.55, M27 objective, or a Leica SP5 laser scanning confocal microscope using the LAS AF 2.2 software (objectives used: 20x 0.70 NA HC PL APO multi-immersion, 40x 1.25 NA, 63x 1.4 NA, or 100x 1.43 NA HCX PL APO oil). Images were analyzed and processed using Volocity (Improvision) and Adobe Photoshop CC (Adobe).

### Reagent Availability

All insect stocks generated as a result of this screen that are not publicly available will be shared upon request. A full listing of the Twist-interacting proteins identified in the double interaction screen is presented in Table 2. Yeast plasmids and strains generated in this study are available upon request.

## RESULTS

### A double interaction screen identifies several context-dependent Twist-interacting proteins

To identify Twist-interacting proteins (TIPs) that regulate Twist transcription factor activity, we used a modified yeast double interaction screen first described by Yu and colleagues (Yu *et al*. 1999). This screen employs both the transcription factor of interest (e.g. Twist) and an established enhancer regulated by that transcription factor as bait (Figure 1A). This strategy affords an advantage over the traditional yeast two-hybrid approach, as it identifies proteins that interact with Twist in the context of a Twist-regulated enhancer. Further, this approach permits the re-screening of identified candidate proteins to determine specificity of the interaction across a variety of Twist-regulated enhancers. Finally, this screening strategy also allows for flexibility in identifying spatially and/or temporally regulated TIPs, by utilizing different cDNA expression libraries. Given our interest in embryonic mesoderm development, our screen utilized a cDNA expression library prepared from early *Drosophila* embryos (Yu *et al*. 1999).

**Figure 1.**
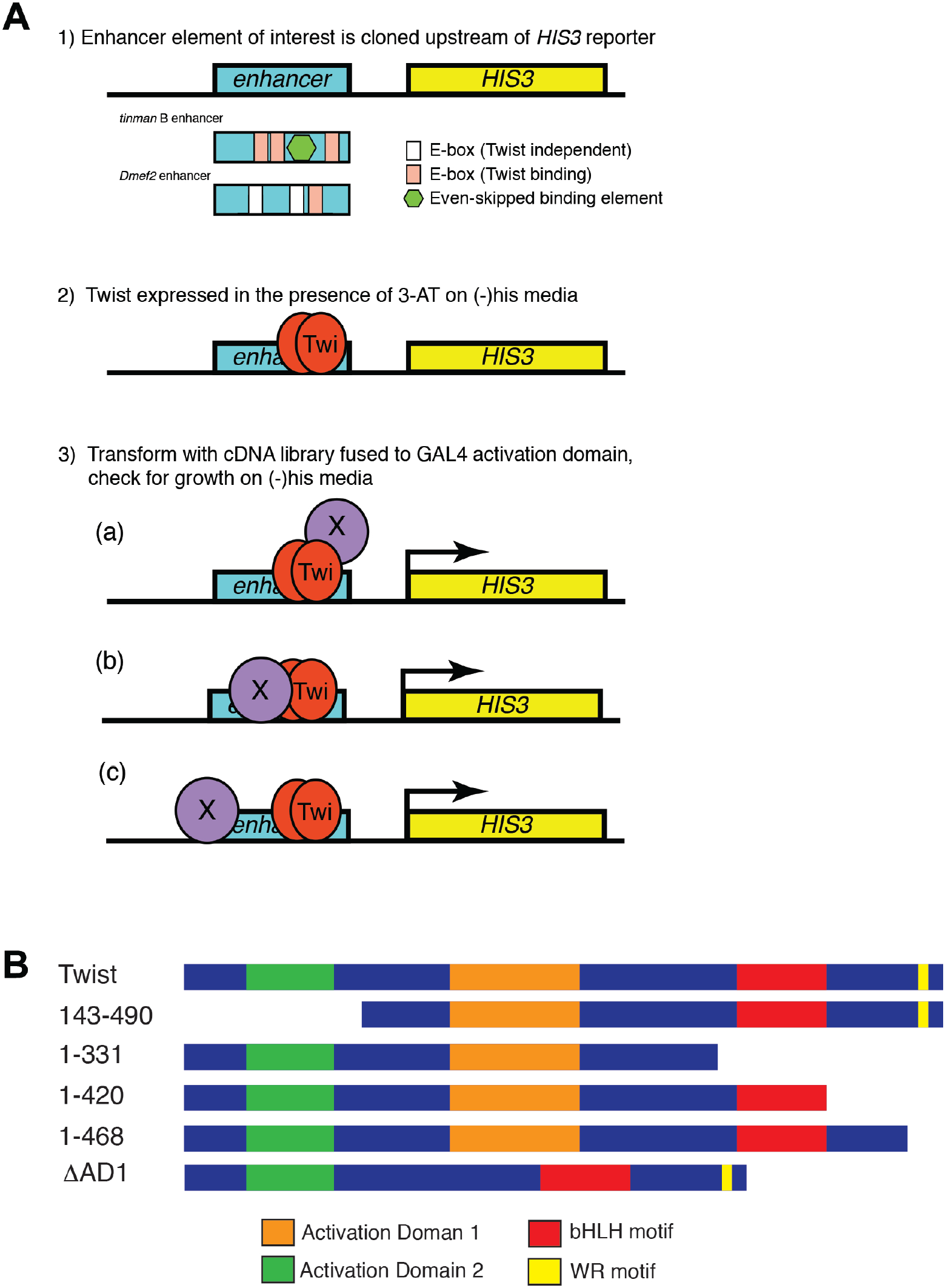
A double interaction screen to identify Twist-interacting proteins (TIPs). (**A**) Schematic of double interaction screen strategy. (1) Enhancer of interest is cloned upstream of the HIS3 reporter cassette in the pACT2 vector and stably transformed into *S. cerevisiae*. For this screen, either a 200 bp fragment containing the Twist-regulated enhancer of the *Dmef2* gene or a 374 bp fragment containing the *tinman* B enhancer element is used. Enhancer-HIS cassette-containing yeast cells are transformed with a Twist expression vector and treated with 3-AT to reduce background Twist activation of HIS reporter. (3) cDNA library, consisting of cDNA from 0-6 hour *Drosophila* embryos fused to GAL4 DNA binding domains is transformed into enhancer-HIS cassette, Twist^+^ yeast cells, and growth on HIS^-^media is determined, indicating a Twist-TIP interaction at the enhancer of interest. Three outcomes are possible for identified TIP proteins: (a) TIPs that interact exclusively with Twist, independent of enhancer binding, (b) TIPs that bind the enhancer cooperatively with Twist, or (c) TIPs that bind the enhancer independently of Twist. (**B**) Schematic of full-length and Twist deletion constructs used to determine region of interaction between Twist and identified TIP proteins.

We chose two Twist-dependent enhancers that are regulatory elements of the *Dmef2* and *tinman* genes (Yin *et al*. 1997; Cripps *et al*. 1998). These two elements were chosen based on their high degree of Twist-dependence and extensive molecular characterization. The endogenous 175bp *Dmef2* enhancer used in our screen is positioned 2 kilobases upstream from the transcription start site (Cripps *et al*. 1998). This simple enhancer contains two E-box consensus sequences, only one of which is solely responsive to Twist activity in the embryonic mesoderm at stage 11 (Figure 1A) (Bour *et al*. 1995; Cripps *et al*. 1998) and in the adult muscle progenitors during larval and pupal stages (Cripps *et al*. 1998; Nguyen and Xu 1998). For the *tinman (tin)* enhancer, we used a 374 bp fragment containing the *tinB* enhancer, located within the first intron of the *tin* locus (Yin *et al*. 1997). This enhancer contains three E-box binding sites for Twist in addition to an Even-skipped homeodomain binding sequence (Figure 1A). This enhancer is responsive to Twist early in development but is not responsive to Twist after stage 10 (Yin *et al*. 1997; Jin *et al*. 2013; Lovato *et al*. 2015) (Figure 1A).

For this screen, each enhancer element was cloned upstream of a HIS3 reporter and transformed into *S. cerevisiae* to create the *Dmef2/HIS* and *tin/HIS* strains. An expression plasmid containing Twist was then transfected into each of the *Dmef2/HIS* and *tin/HIS* strains, leading to a background level of HIS activation. This background was reduced using 12.5mM 3-amino triazol, a specific inhibitor of histidine biosynthesis (Yu *et al*. 1999). Finally, a *Drosophila* embryonic cDNA library, which consisted of the Gal4 activation domain fused in frame with individual cDNAs (Yu *et al*. 1999), was then transformed into both strains.

113 colonies were recovered from the *Dmef2*/*HIS* screen, which were further narrowed down to 23 upon applying several criteria such as plasmid dependence and sequence analysis (Table 1). 117 colonies were recovered from the *tin*/*HIS* screen and 37 of these were pursued after using the above criteria (Table 1). Together, these data suggest that the double interaction yeast screen is capable of identifying new cofactors that interact with Twist in an enhancer specific manner.

**Table 1.** Number of recovered cDNAs and their enhancer specificity identified from each successive step of the double interaction screen.

| Screening step | Enhancer tested |  |
| --- | --- | --- |
|  | *DMef2* | *tinman* |
| Yeast colonies picked | 113 | 117 |
| cDNAs recovered | 88 | 110 |
| cDNAs retested positive | 40 | 65 |
| cDNAs retained after sequencing | 23 | 37 |
| Twist dependent | 20 | 17 |

**Table 2.**
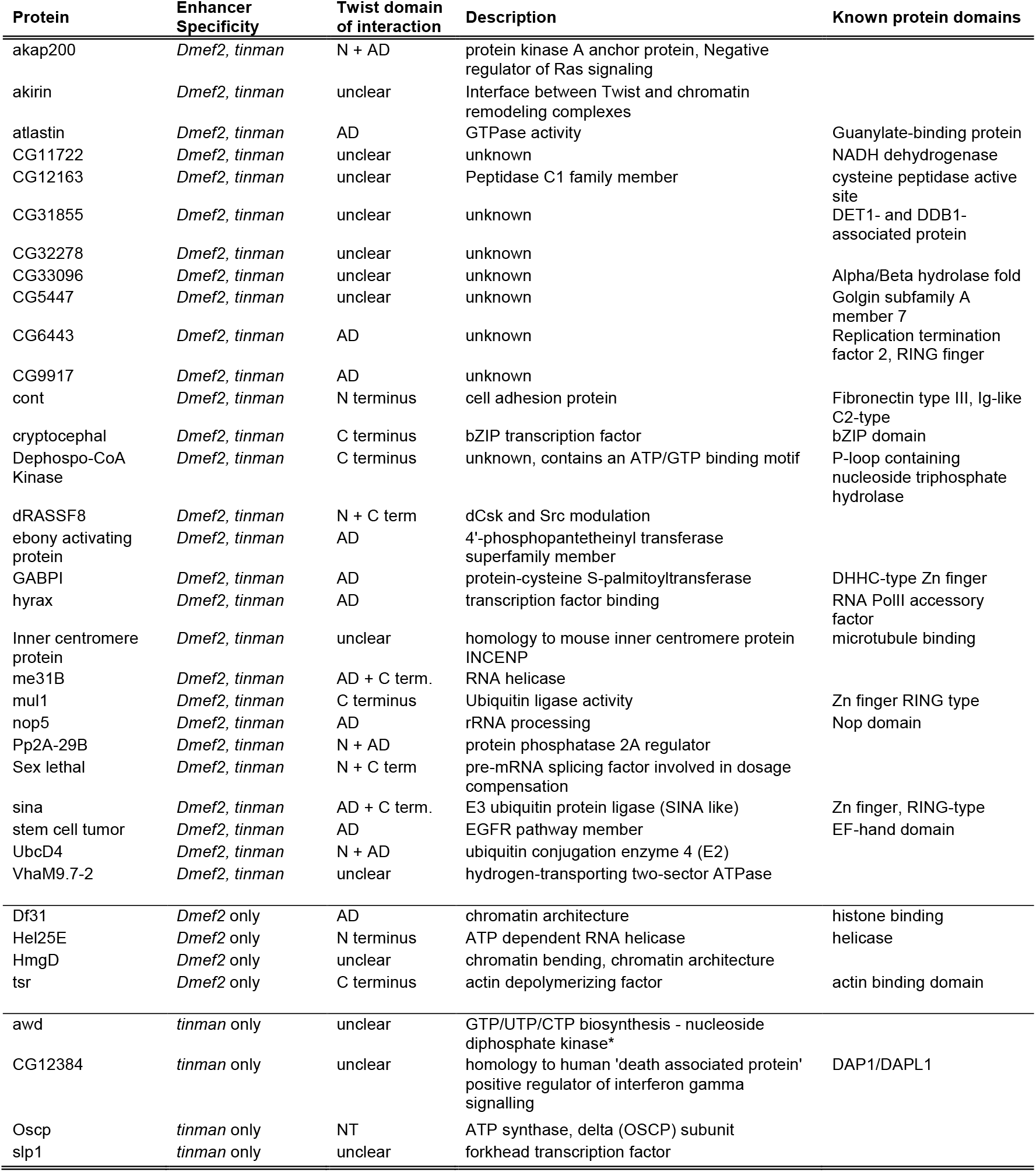
List of Identified Twist-Interacting proteins from double interaction screen and known and/or predicted TIP function.

| Protein | Enhancer Specificity | Twist domain of interaction | Description | Known protein domains |
| --- | --- | --- | --- | --- |
| akap200 | *Dmef2, tinman* | N + AD | protein kinase A anchor protein, Negative regulator of Ras signaling |  |
| akirin | *Dmef2, tinman* | unclear | Interface between Twist and chromatin remodeling complexes |  |
| atlastin | *Dmef2, tinman* | AD | GTPase activity | Guanylate-binding protein |
| CG11722 | *Dmef2, tinman* | unclear | unknown | NADH dehydrogenase |
| CG12163 | *Dmef2, tinman* | unclear | Peptidase C1 family member | cysteine peptidase active site |
| CG31855 | *Dmef2, tinman* | unclear | unknown | DET1- and DDB1-associated protein |
| CG32278 | *Dmef2, tinman* | unclear | unknown |  |
| CG33096 | *Dmef2, tinman* | unclear | unknown | Alpha/Beta hydrolase fold |
| CG5447 | *Dmef2, tinman* | unclear | unknown | Golgin subfamily A member 7 |
| CG6443 | *Dmef2, tinman* | AD | unknown | Replication termination factor 2, RING finger |
| CG9917 | *Dmef2, tinman* | AD | unknown |  |
| cont | *Dmef2, tinman* | N terminus | cell adhesion protein | Fibronectin type III, Ig-like C2-type |
| cryptocephal | *Dmef2, tinman* | C terminus | bZIP transcription factor | bZIP domain |
| Dephospo-CoA Kinase | *Dmef2, tinman* | C terminus | unknown, contains an ATP/GTP binding motif | P-loop containing nucleoside triphosphate hydrolase |
| dRASSF8 | *Dmef2, tinman* | N + C term | dCsk and Src modulation |  |
| ebony activating protein | *Dmef2, tinman* | AD | 4'-phosphopantetheinyl transferase superfamily member |  |
| GABPI | *Dmef2, tinman* | AD | protein-cysteine S-palmitoyltransferase | DHHC-type Zn finger |
| hyrax | *Dmef2, tinman* | AD | transcription factor binding | RNA PolII accessory factor |
| Inner centromere protein | *Dmef2, tinman* | unclear | homology to mouse inner centromere protein INCENP | microtubule binding |
| me31B | *Dmef2, tinman* | AD + C term. | RNA helicase |  |
| mul1 | *Dmef2, tinman* | C terminus | Ubiquitin ligase activity | Zn finger RING type |
| nop5 | *Dmef2, tinman* | AD | rRNA processing | Nop domain |
| Pp2A-29B | *Dmef2, tinman* | N + AD | protein phosphatase 2A regulator |  |
| Sex lethal | *Dmef2, tinman* | N + C term | pre-mRNA splicing factor involved in dosage compensation |  |
| sina | *Dmef2, tinman* | AD + C term. | E3 ubiquitin protein ligase (SINA like) | Zn finger, RING-type |
| stem cell tumor | *Dmef2, tinman* | AD | EGFR pathway member | EF-hand domain |
| UbcD4 | *Dmef2, tinman* | N + AD | ubiquitin conjugation enzyme 4 (E2) |  |
| VhaM9.7-2 | *Dmef2, tinman* | unclear | hydrogen-transporting two-sector ATPase |  |
| Df31 | *Dmef2* only | AD | chromatin architecture | histone binding |
| Hel25E | *Dmef2* only | N terminus | ATP dependent RNA helicase | helicase |
| HmgD | *Dmef2* only | unclear | chromatin bending, chromatin architecture |  |
| tsr | *Dmef2* only | C terminus | actin depolymerizing factor | actin binding domain |
| awd | *tinman* only | unclear | GTP/UTP/CTP biosynthesis - nucleoside diphosphate kinase* |  |
| CG12384 | *tinman* only | unclear | homology to human 'death associated protein' positive regulator of interferon gamma signalling | DAP1/DAPL1 |

|  |  |  |  |
| --- | --- | --- | --- |
| Oscp | *tinman* only | NT | ATP synthase, delta (OSCP) subunit |
| slp1 | *tinman* only | unclear | forkhead transcription factor |

### Secondary screening for Twist interaction and enhancer specificity

From the double interaction screen, 60 positive cDNAs were further analyzed. To eliminate those proteins that bound to either of the enhancers independently of Twist, we tested our 60 putative clones for Twist-dependence. To do this, the candidate cDNAs were transformed into either the *Dmef2/HIS* or *tin/HIS* strain in the absence of the Twist-expressing plasmid. The TIPs were identified by their inability to support growth in the absence of Twist. Those that did support growth under these conditions regulated these enhancers independently of Twist and were eliminated. Using the above criteria, we further eliminated 23 cDNAs that did not exhibit Twist-dependence on either enhancer, furthering shortening our list to 37 TIPs (Table 1). We next investigated whether the TIPs identified were general Twist-interacting partners or enhancer-specific Twist cofactors. To distinguish between these two possibilities, we assayed for the ability of each TIP to activate the reporter of the other enhancer construct. Of the 37 TIPs, 28 tested positively on both enhancers, 5 tested positively only on the *Dmef2* enhancer and 4 tested positively only on the *tin* enhancer. Together, these experiments lead to the recovery of 37 TIPs and demonstrated the sensitivity of our screen to distinguish between general-TIPs and enhancer-specific TIPs (Table 2).

### Identifying Twist protein domain of interaction with TIPs

We next determined the Twist domain(s) necessary for TIP/Twist interactions. We generated Twist constructs that retained the Twist bHLH regions while harboring perturbations in other regions of the protein, as well as constructs that lacked the bHLH domain (Figure 1B). Each of the 37 TIPs were transfected into to the *Dmef2/HIS* or *tin/HIS* yeast strains, along with each of the Twist constructs and scored for their ability to grow in the absence of histidine. We observed that two-thirds of the TIPs (22 of 37) interacted with specific domain(s) of Twist in the presence of a Twist-dependent enhancer (Table 2). The remaining TIPs (15 of 37) did not interact with a specific domain within Twist, suggesting that these factors interact with either multiple domains or the entire Twist protein (Table 3).

**Table 3.** List of Twist-interacting proteins and their preferred Twist domain of interaction

| N-Terminus | N-Terminus and AD2 | AD1 | bHLH<br>Domain | C-Terminus |
| --- | --- | --- | --- | --- |
| Hel25E | Akap200 | Eap | Me31B | Crc |
| Cont | Pp2A-29B | nop5 | Sina | Mul1 |
|  | Dpck | Df31 |  | Tsr |
|  | UbcD4 | AtI |  | Dpck |
|  | GABPI | Hyx |  |  |
|  |  | CG6443 |  |  |
|  |  | CG9917 |  |  |
|  |  | GABPI |  |  |

We next determined if the TIP /Twist interaction(s) followed any specific pattern. We observed that 8 of the TIPs interacted specifically with the glutamate-rich Activation Domain 1 (AD1) domain (Chung *et al*. 1996; Pham *et al*. 1999) (Table 3). This group includes proteins with GTPase activity (Atlastin), proteins that bound to TFs (Hyrax), and others involved in nucleosome assembly (Df31) (Table 3). Five TIPs demonstrated specific interactions with both the N-terminus and Activation Domain 2 of Twist: these included proteins required for protein kinase A anchoring (Akap200), Protein phosphatase 2A regulation (Pp2A-29B), and those with palmitoyltransferase activity (GABPI) (Table 3). Lastly, 4 TIPs interacted specifically with the C-terminus of Twist and play important roles in actin depolymerization (Tsr) and transcriptional regulation (Crc) (Table 3).

### In vivo and in vitro confirmation of screen results

To evaluate the function(s) of the TIPs *in vivo*, we focused on genes for which mutant lines were readily available and included genes with both known and unknown functions. These genes were analyzed based on three attributes: (1) their mRNA expression pattern *in vivo* (based on available modENCODE data), (2) the presence or absence of a muscle development phenotype in the respective mutants, and (3) genetic interactions between the mutant locus and *twist* during embryonic muscle development.

### Genetic interaction of twist with putative Twist-interacting proteins loci in Drosophila embryos

Eleven TIPs identified in our screen were tested for genetic interactions with *twist* by examining the muscle phenotypes in *TIPlocus;twist* double heterozygous mutant embryos. We found that 8 of the 11 *TIP locus;twist* double heterozygotes showed muscle patterning defects (Figure 2 and Table 4). The muscle phenotypes in *TIP locus;twist* double heterozygous embryos ranged from mild to severe defects, consisting of missing muscles, misattached muscles, and misshapen muscles (Figure 2). Most muscle phenotypes were observed in the dorsal and lateral muscle groups [e.g. lateral transverse (LT) muscles 1-4 and dorsal acute (DA) muscle 1], while occasional defects were observed in the ventral muscles [e.g ventral acute (VA) muscles 1,2]. All the observed muscle phenotypes were reminiscent of previously reported phenotypes of embryos that had misregulated or disrupted Twist activity (Wong *et al*. 2008; Nowak *et al*. 2012), or abnormal levels of *Dmef2* expression (Gunthorpe *et al*. 1999; Nowak *et al*. 2012). These data are indicative of genetic interactions between *twist* and the TIP loci.

**Figure 2.**
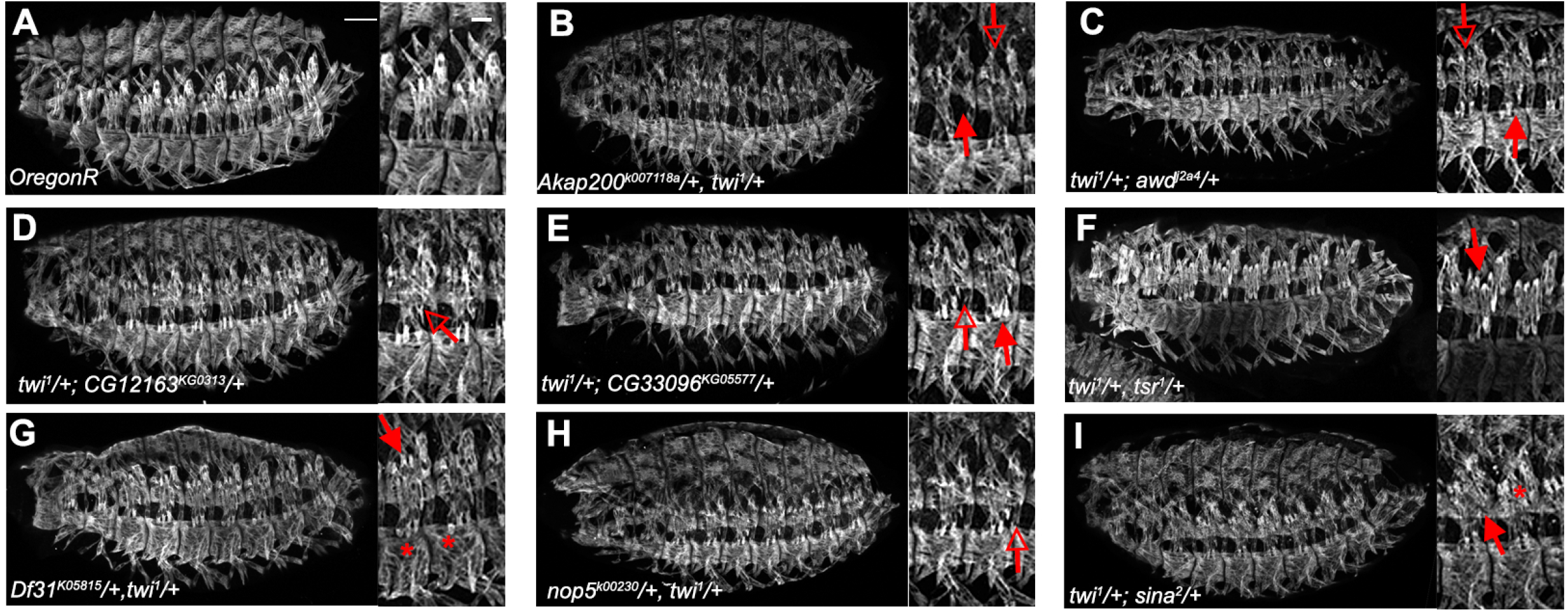
Genetic interactions between *twist* and genes encoding Twist interacting proteins during embryonic myogenesis. (**A-I**) Lateral views of stage 16 wild-type (*OregonR*) and embryos heterozygous for both *twist* and selected genes encoding Twist interacting proteins. All embryos stained to reveal somatic musculature with anti-Myosin heavy chain antibodies. Double heterozygous embryos have defects in general muscle pattering, suggesting a genetic interaction between *twist* and genes encoding TIPs uncovered in double interaction screen. Example phenotypes, such as attachment defects (open arrows), missing or abnormal numbers of muscles (closed arrows), or abnormal muscle morphologies (asterisk) are indicated in inset panels. Scale bar in whole embryo photos = 25 microns. Inset scale bar (all panels) = 10 microns.

**Table 4.**
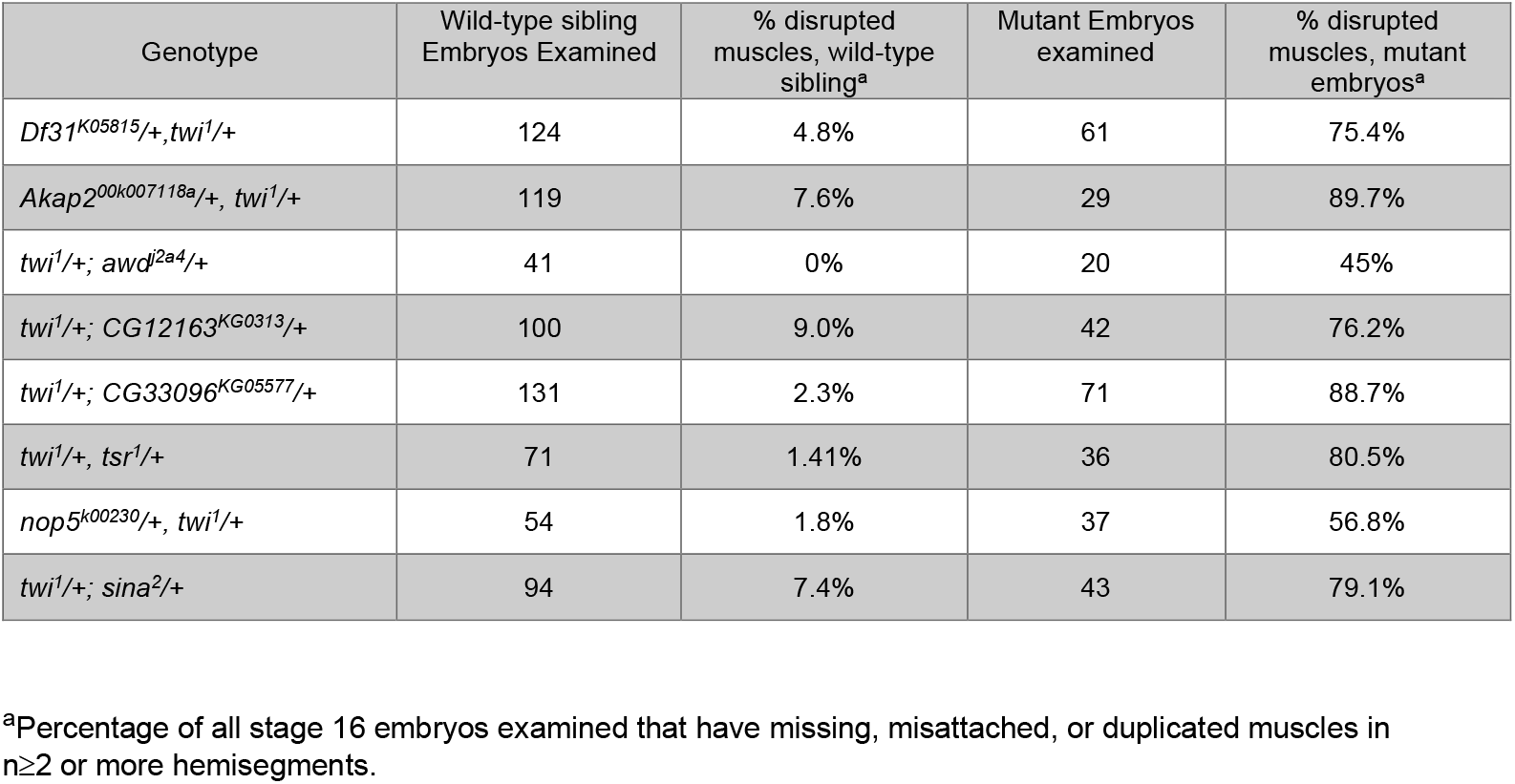
Genetic interactions between identified Twist-interacting loci and *twist*.

### Phenotypes of TIP mutant embryos

Given that our screening methodology identified TIPs in the context of two enhancers critical for both skeletal and cardiac muscle development, we predicted that embryos carrying mutations in these genes would show defects in muscle development. To test this, we examined the muscle phenotype(s) in homozygous mutant embryos for the particular TIP and/or in embryos carrying the mutant allele in *trans* to a deficiency that covered that TIP-encoding locus (Figure 3). Among the TIPs examined, we observed a range of phenotypes from mild to severe (Table 5 and Figure 3). TIP mutant embryos exhibited misattached, misshapen, and/or abnormally attached muscles, particularly in the LT and DA1 muscles (Figure 3). Together, these data suggest that a number of identified TIPs are required for muscle patterning and development.

**Figure 3.**
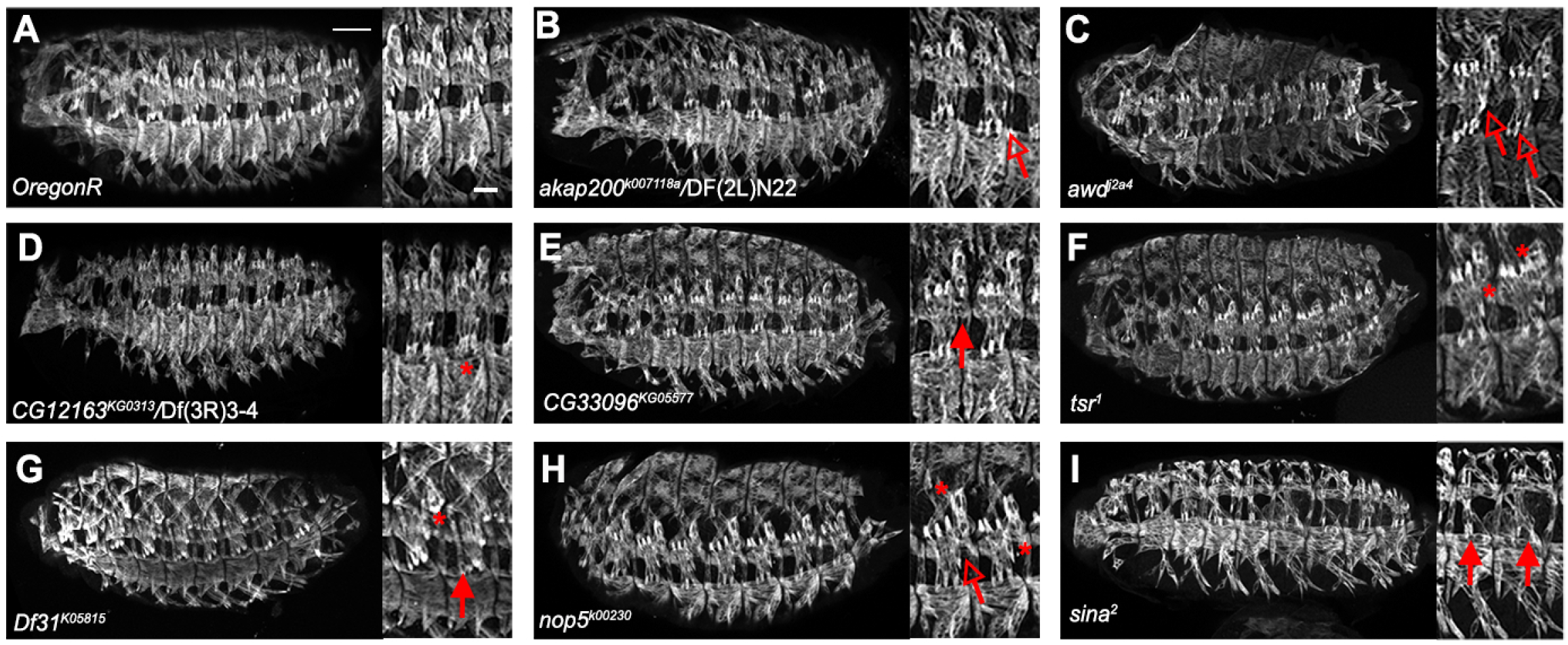
Muscle phenotypes in embryos carrying homozygous mutations in Twist interacting proteins during embryonic myogenesis. (**A-I**) Lateral views of stage 16 wild-type (*OregonR*) and embryos heterozygous for mutations in genes encoding Twist interacting proteins and corresponding chromosomal deficiencies. All embryos stained to reveal somatic musculature with anti-Myosin heavy chain antibodies. In selected mutations of Twist-interacting loci, muscle patterning and morphogenesis aberrations can be observed, ranging from relatively weak to severe. Example phenotypes, such as attachment defects (open arrows), missing or abnormal numbers of muscles (closed arrows), or abnormal muscle morphologies (asterisk) are indicated in inset panels. Scale bar in whole embryo photos = 25 microns. Inset scale bar (all panels) = 10 microns.

**Table 5.**
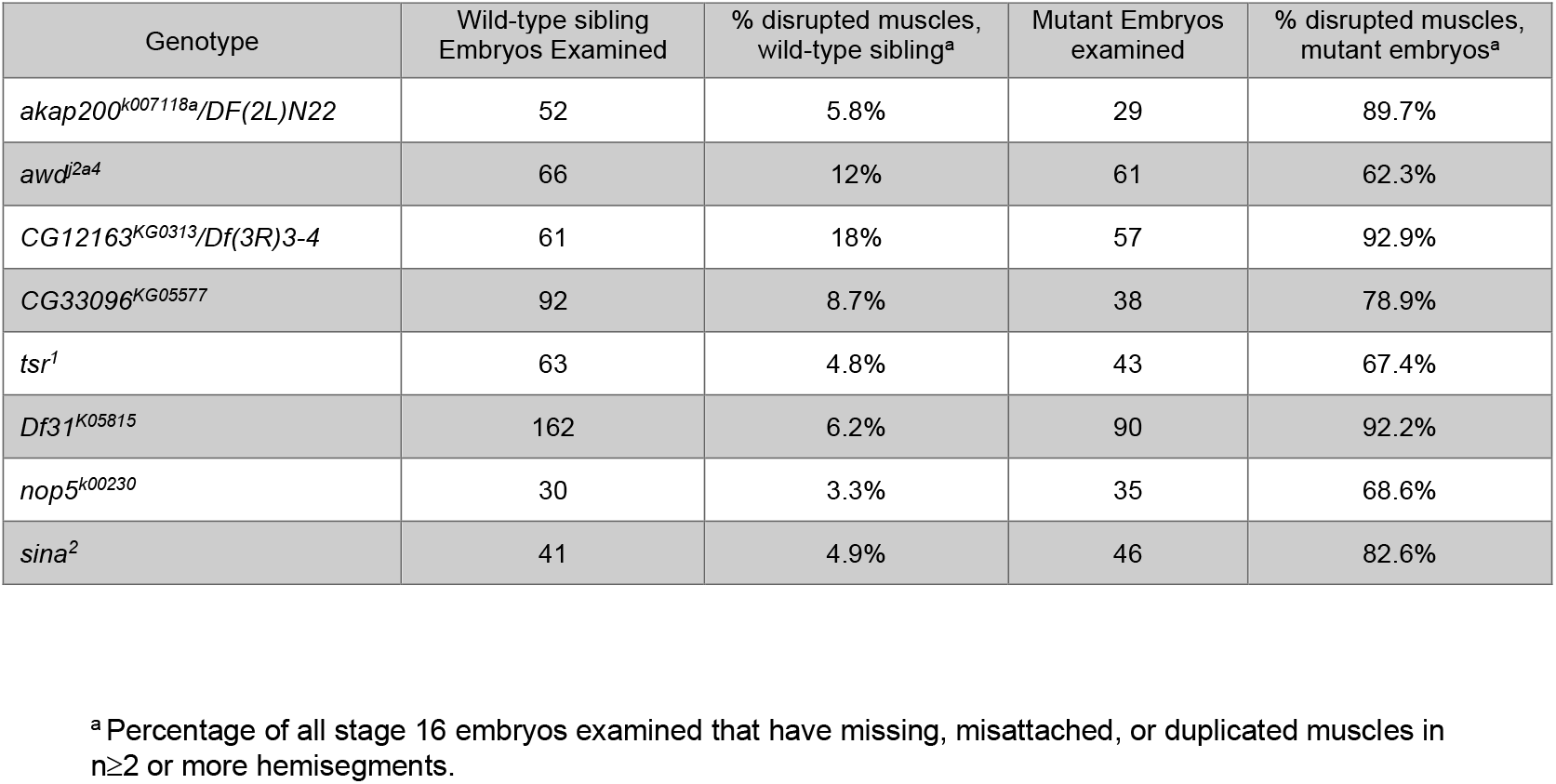
Prevalence of muscle disruptions observed in identified *twist-*interacting homozygotes

### Double interaction screen suggests twinstar as a regulator of embryonic muscle development

The regulation of actin filament organization and structure is a key process throughout myogenesis. Formation of actin structure(s) and their subsequent remodeling occurs throughout myogenesis, including the formation and resolution of F-actin foci during myoblast fusion, formation of tendon/muscle attachment sites, and during assembly and maturation of sarcomeres [reviewed in (Ono 2010; Valdivia *et al*. 2017; Deng *et al*. 2017)]. Moreover, actin and actin-binding proteins localize to the nucleus where they participate in processes such as chromatin remodeling and gene expression (Miralles and Visa 2006; Miyamoto and Gurdon 2012). In vertebrates, actin and actin-binding proteins have been identified as components of transcriptional machineries in muscle nuclei (Favot *et al*. 2005; Miralles and Visa 2006). However, similar roles for the above proteins in *Drosophila* muscles are yet to be identified. Actin depolymerizing factors (ADFs), a subset of actin-binding proteins that sever actin filaments and accelerate filament turnover, play diverse roles throughout development [reviewed in (Kanellos and Frame 2016)]. However, how ADFs impact actin structure(s) during myogenesis remains unknown (Rochlin *et al*. 2010).

Twinstar (Tsr), a member of ADF/Cofilin family of proteins (Edwards *et al*. 1994; Gunsalus *et al*. 1995), was found to interact with the C-terminus of Twist at the *Dmef2* enhancer in our yeast assay. Tsr has been implicated in a number of actin-dependent processes during *Drosophila* development, including cytokinesis in mitotic (larval neuroblast) and meiotic (larval testis) cells, ovary development and oogenesis, protrusion of lamellipodia during border cell (BC) migration, and in the regulation of epithelial integrity of the *Drosophila* wing (Gunsalus *et al*. 1995; Chen *et al*. 2001; Zhang *et al*. 2011; Ko *et al*. 2016). In each of the above contexts, F-actin accumulates aberrantly in *tsr* mutants, indicating that Tsr plays a conserved role in regulating actin filament length, structure, and turnover during these morphogenetic events. Previous work from our lab has focused on the role of actin regulators, primarily actin polymerizing factors, during embryonic muscle development (Richardson *et al*. 2007; Nowak *et al*. 2009). These actin polymerizing factors play important roles in the formation of actin structures throughout myogenesis. Given that Tsr interacts with Twist solely at the somatic musculature enhancer i.e. *Dmef2*, and a role for ADFs/Tsr in *Drosophila* embryonic myogenesis has not yet been described, we chose to further study Tsr and its role in embryonic myogenesis.

### Twinstar localization

We first examined the localization of Tsr at different embryonic stages using a Tsr::GFP protein trap (Morin *et al*. 2001; Kelso *et al*. 2004; Quiñones-Coello *et al*. 2007; Buszczak *et al*. 2007). *tsr* is expressed throughout the embryo at stages 5, 8, 13, and 16 of development (Figures 4A-E). The expression of Tsr coincides with key events in myogenesis, including gastrulation characterized by the expression of *twist* and *Dmef2* (stages 5-8, 2-4 hours AEL), myoblast fusion (stages 12-15, 7-14 hours AEL), myotendinous junction formation and maturation (stages 14-17,10-22 hours AEL), myonuclear movement (stages 14-17, 10-22 hours AEL) and sarcomere assembly (stage 17, 16-22 hours AEL), suggesting that Tsr is involved in mesoderm specification and development. We next determined Tsr’s subcellular localization at stage 16, when a muscle fiber is fully formed, and the nuclei are easily observed (Figure 4). Analysis of fixed embryos revealed that Tsr localizes strictly to the cytoplasm with no obvious subcellular accumulation within the muscle fiber. However, upon observing live embryos, Tsr was found to localize to both the cytoplasmic and nuclear compartments of muscles (Figure 4F). Together, these data indicate that Tsr is expressed in the embryonic musculature and localizes to both the cytoplasm and the nuclei.

**Figure 4.**
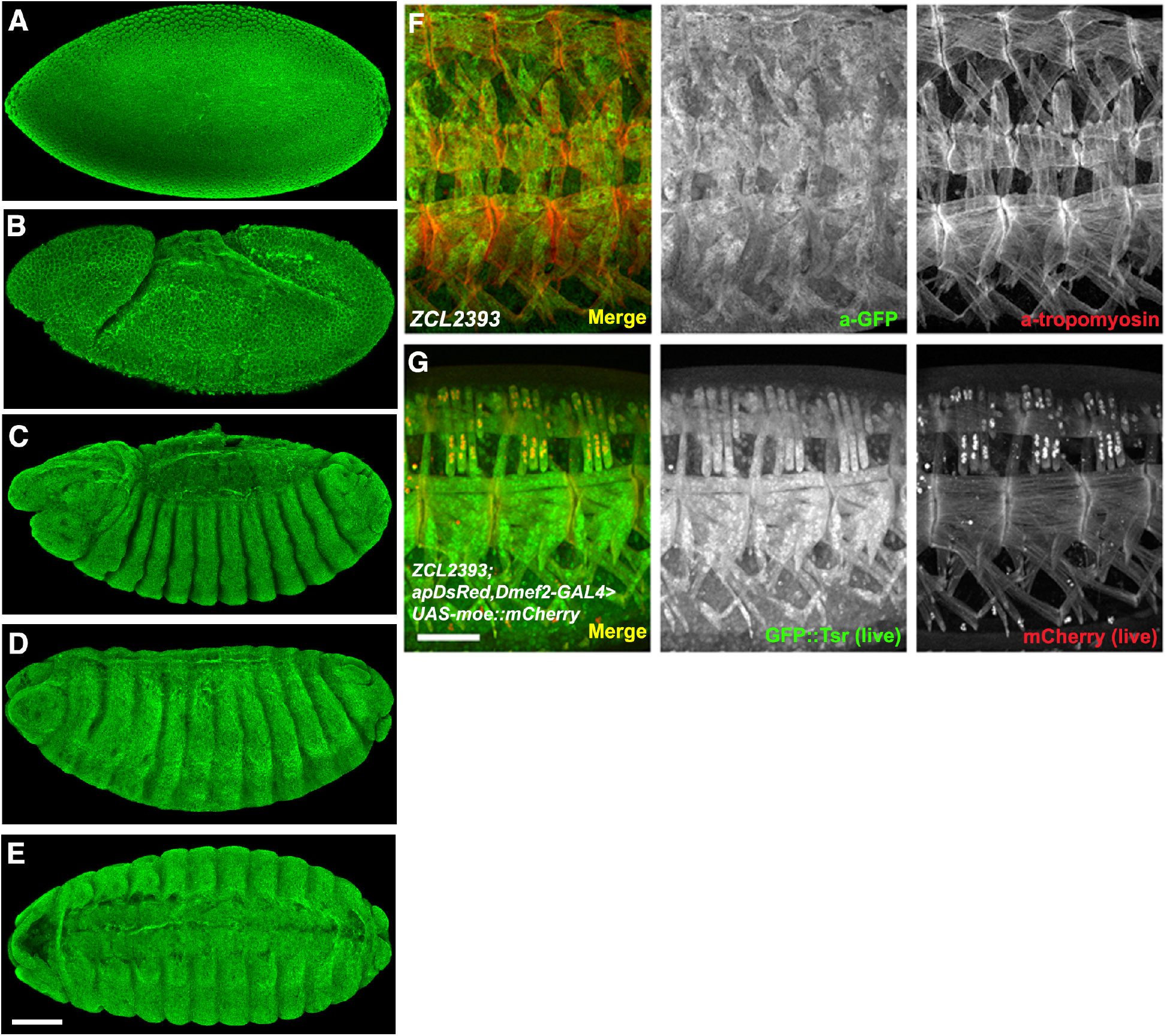
Twinstar is expressed throughout development and in body wall muscles of the *Drosophila* embryo. (**A-E**) Maximum intensity projection of *ZCL2393 (GFP-tsr)* embryos at stage 5 (**A**), stage 8 (**B**), stage 13 (**C**) and stage 16 (**D**, lateral view and **E**, ventral view) labeled with an antibody against GFP (green) to label GFP::Tsr. Tsr is expressed ubiquitously in the embryo. (**F**) Maximum intensity projections of a fixed stage 16 embryo labeled with an antibody against GFP (green) to label GFP::Tsr and Tropomyosin (red) to visualize the embryonic body wall muscle. Three representative hemisegments are shown. (**G**) Endogenous GFP::Tsr in a live embryo without antibody staining. A live reporter for muscle (*apRed, Dmef2-Gal4>UAS-moe::mCherry*), which labels muscle actin and the nuclei of the lateral transverse (LT) muscles, was used to identify the body wall muscle. Bar, 50 μm.

### Twinstar is required for muscle development in Drosophila

To determine if Tsr is required for muscle development and differentiation, we obtained two well-characterized *tsr* mutant alleles (Gunsalus *et al*. 1995). *tsr^1^* is a hypomorphic allele generated by the insertion of a lacZ enhancer trap/P-element in the 5’ UTR of *tsr* that results in approximately 80% reduction in *tsr* transcript levels (Gunsalus *et al*. 1995). *tsr^N96A^* (Ng and Luo 2004) is a null allele generated from the imprecise excision of a P-element from the *tsr* locus (Figure 5A).

**Figure 5.**
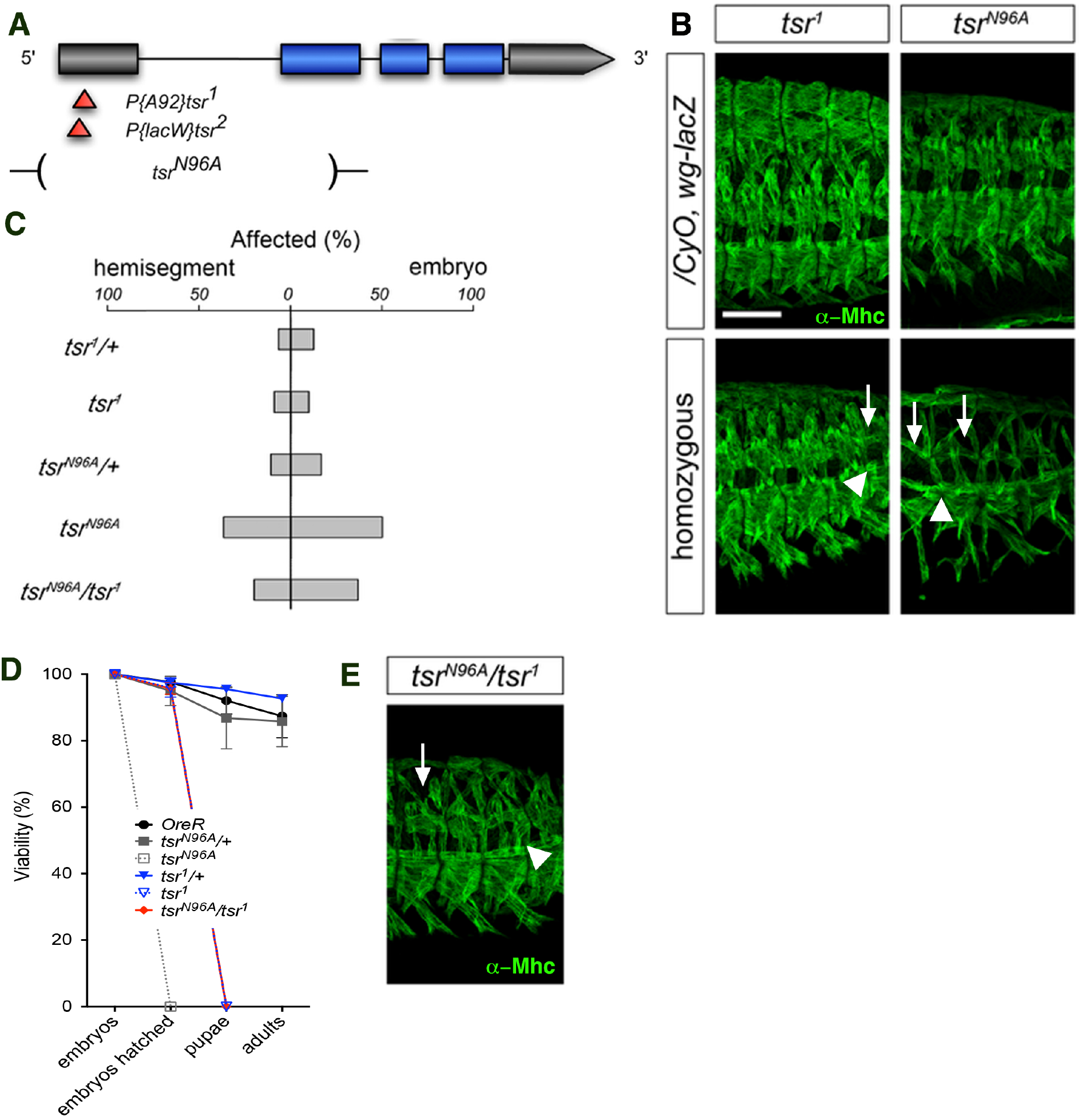
*twinstar (tsr)* mutants have muscle phenotypes. (**A**) Schematic diagram of the *tsr* locus in *Drosophila*. Alleles used in this study are indicated below the locus. Grey bars, untranslated regions. Blue bars, exons. Red and orange triangles, P-element insertions. Deletions that remove portions of the *tsr* locus are schematized by a gap in the chromosome bracketed by parentheses. (**B**) Maximum intensity projection of a stage 16 embryo labeled with an antibody against Myosin heavy chain (green) to visualize the embryonic body wall muscle. Three representative hemisegments are shown for each genotype. Arrows indicate missing muscles. Arrowheads indicated misattached muscles. (**C**) Quantification of percentage of affected hemisegments (left) and embryos (right) of each genotype indicated. n= 20 embryos, 100 hemisegments. (**D**) Viability of the indicated genotypes. n=100. (**E**) Maximum intensity projection of a stage 16 embryo labeled with an antibody against Myosin heavy chain (green) to visualize the embryonic body wall muscle. Three representative hemisegments are shown for each genotype. Arrows indicate missing muscles. Arrowheads indicated misattached muscles. Scale bar, 50 μm.

Examination of *tsr^N96A^* mutant embryos revealed a number of muscle defects including missing and misattached muscles (Figures 5B and 5C). These embryos failed to hatch, indicating that Tsr is essential for organismal viability (Figure 5D). The *tsr^1^* mutant embryos had fewer muscle defects, consistent with *tsr^1^* being a hypomorphic allele (Figures 5B and 5C). While these embryos hatched into larvae, the larvae failed to develop past the first instar (L1) stage (Figure 5D). *tsr^N96A^/tsr^1^* transheterozygous embryos displayed similar defects in embryonic muscle patterning as seen in each of the single *tsr* mutants, i.e. *tsr^96A^* and *tsr^1^*, and a viability profile similar to *tsr^1^* mutant embryos (Figures 5D and 5E). Together, these data indicate that Tsr is essential for muscle development and organism viability.

### Twinstar-interacting proteins are required for proper muscle development

The regulation of ADF/cofilin activity by the phosphorylation state of Ser3 has been well-documented (Bamburg 1999). Phosphorylation by members of two kinase families, LIM domain-containing kinases (LIMK1/2, *Drosophila* LIMK1) and testis-specific protein kinases [TESK1/2, *Drosophila* Center divider (Cdi)], inhibits ADF/cofilin activity, while dephosphorylation by two phosphatases, Slingshot (Ssh) and Chronophin (CG5567 in *Drosophila*), activates ADF/cofilin family members (Morgan *et al*. 1993; Agnew *et al*. 1995; Bamburg 1999; Wang *et al*. 2007; Bravo-Cordero *et al*. 2011). In addition, AIP1 [*Drosophila* Flare (Flr)] synergizes with ADF/cofilin to drive actin dynamic processes (Bamburg 1999). We next asked if Tsr’s role in muscle development was regulated through the same sets of regulators.

We first determined if these genes were expressed in the muscle and then where their proteins were localized. We turned to surveying existing literature, published *in situ* databases, and available protein traps for Tsr’s regulators and its interacting partners. LIMK1 had been observed to be ubiquitously expressed in the embryonic muscle throughout development, while CG5567, though not detected in the body wall muscle, is expressed in pharyngeal and visceral muscle types (Tomancak *et al*. 2002; 2007). Similarly, Flr had previously been reported to localize to the embryonic body wall musculature (Tomancak *et al*. 2002; 2007). Using the *GFP::Flr* protein trap (Morin *et al*. 2001; Kelso *et al*. 2004; Quiñones-Coello *et al*. 2007; Buszczak *et al*. 2007), we confirmed Flr’s localization in the body wall muscles, where it was enriched at the MTJs (Figure 6A). These data suggest that along with Tsr, Tsr’s regulators also localize to the embryonic musculature.

**Figure 6.**
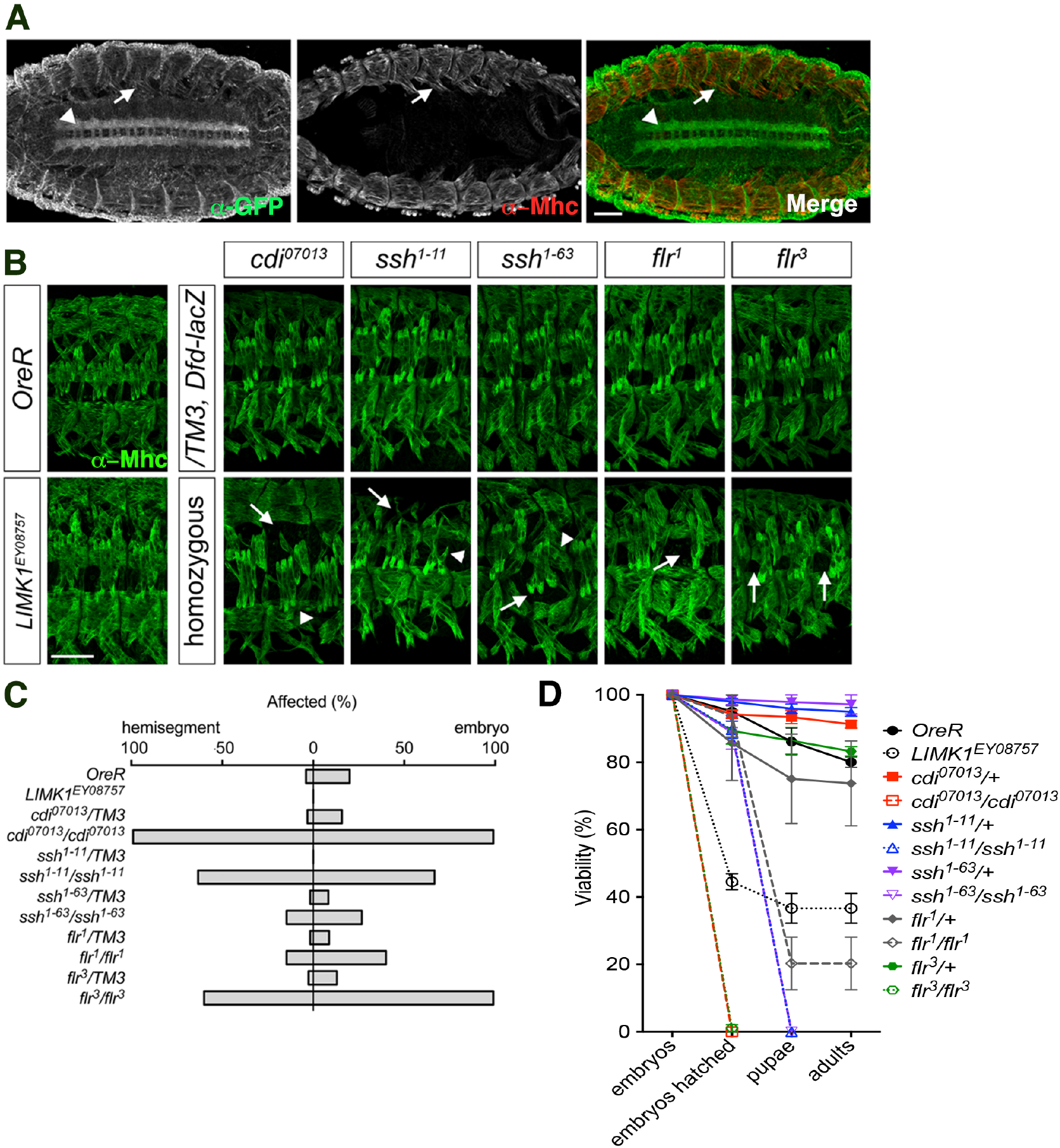
*twinstar’s* regulators are also required for muscle development. (**A**) Maximum intensity projection of a stage 16 embryo labeled with an antibody against GFP (green,α-GFP), to label GFP::Flr, and Myosin heavy chain (red, α-Mhc), to label body wall muscles. Arrows indicate muscle expression. Arrowheads indicate ventral nerve cord expression. (**B**) Maximum intensity projection of stage 16 embryos labeled with an antibody against Myosin heavy chain (green) to visualize the embryonic body wall muscle. Three representative hemisegments are shown for each genotype. Arrows indicate missing muscles. Arrowheads indicate misattached muscles. (**C**) Quantification of percentage of the percentage of affected hemisegments (left) and embryos (right) of the genotypes indicated. n=20 embryos, 100hemisegments. (**D**) Viability of the indicated genotypes.n=100. Scale bar, 50 μm.

To understand the role(s) of Tsr’s regulators in embryonic myogenesis, we obtained mutant alleles for *LIMK1, cdi, ssh*, and *flr*. Embryos homozygous for *LIMK1^EY08757^* appeared wild type (Figures 6B and 6C), concomitant with the likelihood of it being a hypomorphic allele. *LIMK1^EY08757^* flies were homozygous viable and fertile though there was an overall decrease in fitness (Figure 6D). *cdi^07013^* embryos, which contain an intronic P-element insertion (Spradling *et al*. 1999), had a number of muscle defects and failed to hatch into larvae (Figures 6B and 6C). Similarly, *ssh^1-11^* and *ssh^1^*^-63^ mutant embryos had both missing and misattached muscles, and these embryos failed to hatch (Figures 6B–6D). Two EMS-generated alleles of *flr*, *flr^1^* and *flr^3^*, also displayed general muscle patterning defects. *flr^3^* mutant embryos, had a more penetrant muscle phenotype, and subsequently failed to hatch, while the majority of *flr^1^* mutants died during larval development (Figures 6B–6D). Together, these data indicate that the Tsr regulators Cdi, Ssh and Flr are essential for *Drosophila* development and are required for proper muscle morphogenesis in the embryo.

## DISCUSSION

Twist regulates numerous steps during mesoderm development and muscle formation, which suggest that its activity is tightly regulated. While each step relies on differences in Twist activity levels (Wong *et al*. 2014), it is possible that these differences are achieved by not only varying *twist* expression levels through transcription, but also regulating Twist post-translationally. To examine putative post-translational regulation of Twist activity, we performed a double interaction screen to identify potential novel Twist Interacting Proteins (TIPs) that could modulate its activity. Using two well-characterized Twist-regulated enhancers critical for somatic and cardiac muscle development, our screening method uncovered 37 TIPs that interact directly with Twist at either the *Dmef2* enhancer, the *tin* B enhancer, or at both. We confirmed that a subset of these identified TIPs genetically interacted with Twist and that TIP mutant embryos showed muscle patterning phenotypes. The TIPs identified from the screen function in a range of processes, including actin remodeling, transcriptional regulation, chromatin remodeling, and ubiquitination, all which appear to be required for myogenesis (Table 2). This range of different functions suggest that there are multiple modalities for modulating, enhancing, or attenuating Twist TF activity. Our results described here, along with previously published studies on another TIP identified through this screen (Nowak *et al*. 2012) suggest that Twist activity is regulated through a variety of direct and indirect proteinprotein interactions during mesodermal development. Together, these data highlight the efficacy and specificity of our screening methodology to identify novel Twist co-factors.

Twist’s interaction(s) with its co-regulators Daughterless and Dorsal have been well documented. We were interested in extending this understanding to the TIPs identified by our screen. While our data have shed insights to the region(s) of Twist that interact with the TIPs (Table 3), an overall predictive schematic for Twist regulation by these partners remains elusive. Moreover, our data indicate that *twist* genetically interacts with a number of TIP-encoding loci, as revealed by our results showing defects in embryonic muscle patterning. Furthermore, a subset of TIP mutant embryos analyzed displayed defects in somatic muscle formation and patterning, further validating our double interaction screen results as *bona fide* TIPs at the *tin* B and *Dmef2* myogenic enhancers. Future studies to determine the function(s) of the identified TIPs and the processes they control will help to better understand how the TIPs work with Twist to mediate Twist’s different functions during muscle development.

Our screen has uncovered a number of TIPs with diverse functions (Table 2). Though our primary interest was in identifying transcriptional co-regulators of Twist, we also uncovered factors whose function(s) suggest post-translational regulation of Twist, either through phosphorylation or ubiquitination. These post-translational mechanisms of Twist regulation have also been identified in in cancer cell progression (Hong *et al*. 2011), as well as during neural crest specification in *Xenopus* (Laursen *et al*. 2007; Lander *et al*. 2013), suggesting that they are not unique to the muscle and could mediate Twist’s role in a variety of tissues. Our screen has also identified a number of TIPs with reported or predicted activities in other biological processes, including RNA metabolism, transcriptional regulation, chromatin structure and remodeling, post-translational modification, and cell-cell signaling, as well as TIPs with unknown structures or functions (Table 2).

We evaluated a subset of the identified TIPs in detail for putative *in vivo* roles during muscle development. This subset included those with diverse cellular functions, including a protein kinase A scaffolding protein (*Akap200*), chromatin assembly and remodeling factors (*akirin* and *Df31*), actin remodeling factor (*tsr*), peptidase activity (*CG12163*), regulators of ubiquitination (*sina*), transcription activator (*slp1*), and an RNA-processing protein (*nop5*). Despite the diverse nature of the TIPs that were recovered in our screen, we note that embryos bearing mutations in each of these genes showed a variety of defects in muscle development, which would be consistent with a putative role as a Twist-interacting protein.

Given that our interest lay in other cofactors known to affect myogenesis, we examined *sloppy paired 1 (slp1)*, which is critical for embryonic muscle development in *Drosophila* (Riechmann *et al*. 1997). Slp1 is a member of the forkhead-family of transcription factors, known to repress *bagpipe* expression in the mesoderm and is necessary, but not sufficient, for the specification of both somatic muscle and cardiac precursors (Lee and Frasch 2000). In our screen, Slp-Twist interactions were restricted to the *tinman* enhancer. This interaction is expected as the *tinman* enhancer used in our screen contains a sequence similar to a Slp1 binding site in the *fushi-tarazu* promoter (Lee and Frasch 2000), while no such sites exist within the *Dmef2* enhancer.

Some of the TIPs identified possess ubiquitin ligase activity. Of these, *seven in absentia* (*sina*) encodes a RING-type E3 ubiquitin ligase that facilitates the transfer of ubiquitin from the E2 ubiquitin-conjugating enzyme to the substrate (Li *et al*. 2002). Our examination of *sina* mutant embryos revealed defects in the embryonic muscle pattern, indicating its possible role in muscle development (Figure 5). These data are consistent with previous data that reported the ubiquitous expression of *sina* in the mesoderm. Moreover, *sina* has been shown to be required for the ubiquitination of the zinc finger transcription factor Tramtrack (Ttk) during the formation of the R7 photoreceptor in the *Drosophila* eye (Li *et al*. 2002; Artero *et al*. 2003). Recent work has demonstrated Trk as a key repressor of founder cell fate in fusion-competent myoblasts during myogenesis (Ciglar *et al*. 2014). Sina was one of the TIPs in our screen that possessed either E2 or E3 ubiquitin ligase activity, suggesting that Twist could be a possible target of ubiquitination. Twist ubiquitination (Lander *et al*. 2013) and phosphorylation (Hong *et al*. 2011; Lander *et al*. 2013) at the WR domain have been identified as key regulatory processes that control Twist’s stability in a variety of developmental processes. However, it remains to be determined if Twist exhibits similar post-translational modifications in the context of the myogenic gene regulatory program.

Our final TIP of interest was A kinase anchor protein 200 (Akap200), which interacts with Twist at both the *Dmef2* and *tinman* enhancers. Akap200 is a kinase anchoring protein, known to tether protein kinase A activity to specific cellular domains (Jackson and Berg 2002; Bonin and Mann 2004). Akap200 is also known to tether kinase A activity to the actin cytoskeleton during axon outgrowth (Huang and Rubin 2000; Jackson and Berg 2002), as well as to negatively regulate Ras activity (Huang and Rubin 2000). Both the actin cytoskeleton and Ras signaling play important roles during muscle development (Carmena *et al*. 2006). *Akap200* mutant embryos showed defects in muscle development (Figures 2 and 3), suggesting a direct role for Akap200 in Twist-mediate myogenic gene expression. Future studies to characterize these genes in detail could aid in understating their role(s) in muscle development and in the regulation of Twist activity.

Actin is critical for embryonic muscle development, as actin is remodeled into different actin structures in the myoblasts and myotubes are required to form a functional muscle fiber, while nuclear actin and actin-binding proteins are components of chromatin and transcription complexes (Richardson *et al*. 2008; Ono 2010; Deng *et al*. 2017). Twinstar (Tsr), a member of the ADF/Cofilin family of proteins, has been shown to regulate actin filament length and structure in embryonic tissues, while its role in the embryonic muscle has not been investigated. We recovered Tsr as a Twist interacting protein at the *Dmef2* enhancer. Examination of *twist;tsr* double heterozygous embryos showed defects in muscle structure, confirming genetic interaction between Twist and Tsr *in vivo*. Similarly, embryos mutant for *tsr* or its regulators show muscle defects, with severity in muscle defects correlating with severity of *tsr* mutation. These data suggest that Tsr and its regulators play important role(s) in the patterning and development of the embryonic musculature. However, future experiments to identify the precise step(s) where Tsr and its regulators function can shed insight to their functions during myogenesis.

How and where Tsr regulates Twist activity is a remaining question from this study. Tsr could interact with Twist directly (i.e. Tsr binding to Twist could function as a cofactor), or indirectly (i.e. Tsr could interact with RNA-pol II) to facilitate Twist-mediated transcription. Recently, Tsr was found to play a role in the heat mediated actin stress response (ASR) pathway (Figard *et al*. 2019). Tsr’s activity was found to increase during ASR, leading to increased F-actin severing in the cytoplasm and an increase in G-actin, which is subsequently transported into the nucleus and forms actin rods (Figard *et al*. 2019). As actin does not possess a classic Nuclear localization sequence (NLS), Tsr, via its NLS (Gunsalus *et al*. 1995), is thought to interact with importins to mediate transport of actin into the nucleus (Dopie *et al.*). In this study, we observe Tsr expression in myonuclei, suggesting a possible nuclear function for Tsr. Tsr, similar to its vertebrate Cofilin counterpart, could be a component of RNA-Pol II transcriptional machineries where in conjunction with actin they could be responsible for transcriptional elongation of coding sequences, by maintaining the monomer state of actin (Obrdlik and Percipalle 2011). Hence, Tsr could indirectly be influencing Twist transcriptional activity through its effects on RNA polymerase. Alternatively, Tsr may directly interact with Twist, and this Tsr interaction with a transcription factor would represent a novel function for the ADF/Cofilin family. We have found that Tsr was found to interact with C-terminus of Twist at the *Dmef2*-enhancer in our screen. This interaction between Twist and Tsr could occur in the nucleus as part of the transcription complex along with actin. Future experiments that probe the interaction between the two proteins in different subcellular compartments would help shed insight to their interactions and other binding partners.

The structure of Twist, its function, and its target genes have been studied across metazoans within multiple developmental contexts. However, how Twist regulates this diverse transcriptional profile and its interactions with co-factors, i.e TIPs, are only now being understood. Our study employing the double interaction screening strategy has successfully identified a host of new Twist-interacting proteins at two well-studied Twist-regulated enhancers required for the regulation of the somatic and cardiac muscle lineages in *Drosophila*. It remains to be determined whether the roles of this cast of cofactors is unique to these particular enhancers or is part of a more universal method of Twist regulation in other developmental contexts and organisms. Finally, identification of a role for Tsr in embryonic muscle development validates our screen for identifying TIPs and presents an exciting new avenue to study Cofilin regulation, the interactions between actin cytoskeletal regulators and transcription factors such as Twist during myogenesis.

## ACKNOWLEDGEMENTS

The authors wish to thank L. Pick and R. Carthew for the gift of reagents, David Soffar for technical assistance, as well as members of the Baylies and Nowak laboratories for critical comments on the finished manuscript. Stocks obtained from the Bloomington *Drosophila* Stock Center (NIH P40OD018537) were used in this study. This work was supported by NIH GM102826, NSF DBI-1229237, and American Heart Association AIREA Award 33960109 to SJN, a National Cancer Institute [P30 CA 008748] core grant to MSKCC, and NIH AR068128, NIH AR067361, and MDA Research Development Grant MDA4153 to MKB.

---

**Figure 1. A double interaction screen to identify Twist-interacting proteins (TIPs)**.

**Figure 2. Genetic interactions between *twist* and genes encoding Twist interacting proteins during embryonic myogenesis**.

**Figure 3. Muscle phenotypes in embryos carrying homozygous mutations in Twist interacting proteins during embryonic myogenesis**.

**Figure 4: Twinstar (Tsr) is expressed in the body wall muscles of the *Drosophila* embryo**

**Figure 5: *twinstar (tsr)* mutants have muscle phenotypes**

**Figure 6: Regulators of *twinstar are* also required for muscle development**

